# Dynamical structure-function correlations provide robust and generalizable signatures of consciousness in humans

**DOI:** 10.1101/2023.12.19.572402

**Authors:** Pablo Castro, Andrea Luppi, Enzo Tagliazucchi, Yonatan S-Perl, Lorina Naci, Adrian M. Owen, Jacobo D. Sitt, Alain Destexhe, Rodrigo Cofré

## Abstract

Resting-state functional magnetic resonance imaging evolves through a repertoire of functional connectivity patterns which might reflect ongoing cognition, as well as the contents of conscious awareness. We investigated whether the dynamic exploration of these states can provide robust and generalizable markers for the state of consciousness in human participants, across loss of consciousness induced by general anaesthesia or slow wave sleep. By clustering transient states of functional connectivity, we demonstrated that brain activity during unconsciousness is dominated by a recurrent pattern primarily mediated by structural connectivity and with a reduced capacity to transition to other patterns. Our results provide evidence supporting the pronounced differences between conscious and unconscious brain states in terms of whole-brain dynamics; in particular, the maintenance of rich brain dynamics measured by entropy is a critical aspect of conscious awareness. Collectively, our results may have significant implications for our understanding of consciousness and the neural basis of human awareness, as well as for the discovery of robust signatures of consciousness that are generalizable among different brain conditions.

## Introduction

The identification of reliable markers for determining the presence or absence of human consciousness from brain signals remains a major unresolved problem in neuroscience (Sarasso et al. 2015; Demertzi et al. 2019; Rankaduwa and Owen 2023; Casali et al. 2013; Dehaene and Changeux 2011; Casarotto et al. 2016; Bodart et al. 2017; Gosseries et al. 2014; Zilio et al. 2023). The spontaneous dynamics of the brain can be measured using resting-state functional magnetic resonance imaging (rs-fMRI) (B. B. Biswal 2012; B. Biswal et al. 1995; Allen et al. 2014; Cabral et al. 2017; Cornblath et al. 2020), consisting of the recording of functional magnetic resonance imaging while no task is explicitly performed (i.e. during rest). Using this approach, it has been shown that the brain exhibits dynamic transitions between various transient configurations, each serving distinct cognitive functions (Deco, Jirsa, and McIntosh 2013; Shine et al. 2016; Lurie et al. 2020; Fox et al. 2006; Vincent et al. 2006; Damoiseaux et al. 2006; De Luca et al. 2005; Raichle et al. 2001; Calhoun et al. 2014; Preti, Bolton, and Van De Ville 2017; Baker et al. 2014; Deco, Jirsa, and McIntosh 2011). The nature of signals during the resting state suggest that they might reflect complex neural processes related to ongoing cognition and consciousness, prompting the search for signatures of consciousness in the dynamics of spontaneous brain activity (Demertzi et al. 2019; Luppi et al. 2019; Luppi, Carhart-Harris, et al. 2021; Barttfeld, Uhrig, et al. 2015; Huang et al. 2020; Hudetz, Liu, and Pillay 2015; Mortaheb et al. 2022). For instance, differences in the sequences and prevalence of transient brain states measured in macaques can distinguish between wakefulness and general anesthesia induced by propofol, ketamine and sevoflurane (Barttfeld, Uhrig, et al. 2015; Uhrig et al. 2018), a finding that was replicated in human individuals suffering from disorders of consciousness (Luppi, Golkowski, et al. 2021; Huang et al. 2020; Demertzi et al. 2019).

One of the major obstacles behind the aforementioned challenge consists of determining robust and generalizable markers of consciousness (Sarasso et al. 2021; Bai, Lin, and Ziemann 2021; Massimini et al. 2005). In this context, robust indicates markers which do not require fine-tuning of parameters for each different dataset where they are applied; moreover, robustness also stands for the independence of the consciousness state of the individuals. Generalizability refers to the capacity to successfully detect consciousness markers in different datasets obtained using heterogeneous acquisition parameters and under potentially divergent experimental conditions. Currently, few papers explicitly addressed these two requirements, especially in combination. This represents an important obstacle, both for the study of fundamental principles underlying consciousness, and for the translation to the clinics.

Here we focused on determining markers of consciousness based on the dynamics of functional connectivity measured using rs-fMRI in human participants, contrasting the awake state with general anaesthesia (intravenous propofol) and deep sleep, two different brain states characterised by diminished or absent conscious awareness, and corresponding to two datasets acquired by different research groups using different parameters of data acquisition. Our results provide evidence that consciousness exhibits specific features that are grounded in the temporal dynamics of ongoing brain activity, which are common to different brain states, and robust against changes in experimental acquisition protocols.

## Mathematical Methods

Before entering into the details, we describe here the general idea of the methodology for which a summary can be seen in Figure 1. We first consider in both datasets blood-oxygen-level-dependent (BOLD) signals of healthy volunteers. For each participant we computed the Z-scored BOLD time series (Fig 1A). Secondly, we compute the Hilbert transform of the said signals to extract the phase of the signal for each region of interest (ROI) at each repetition time (TR) (Fig 1B & 1C). We next compare every ROI pair of these phases at every TR to determine brain regions’ synchronizations and anti-synchronizations evolving through time (Fig 1D).

**Figure 1.**
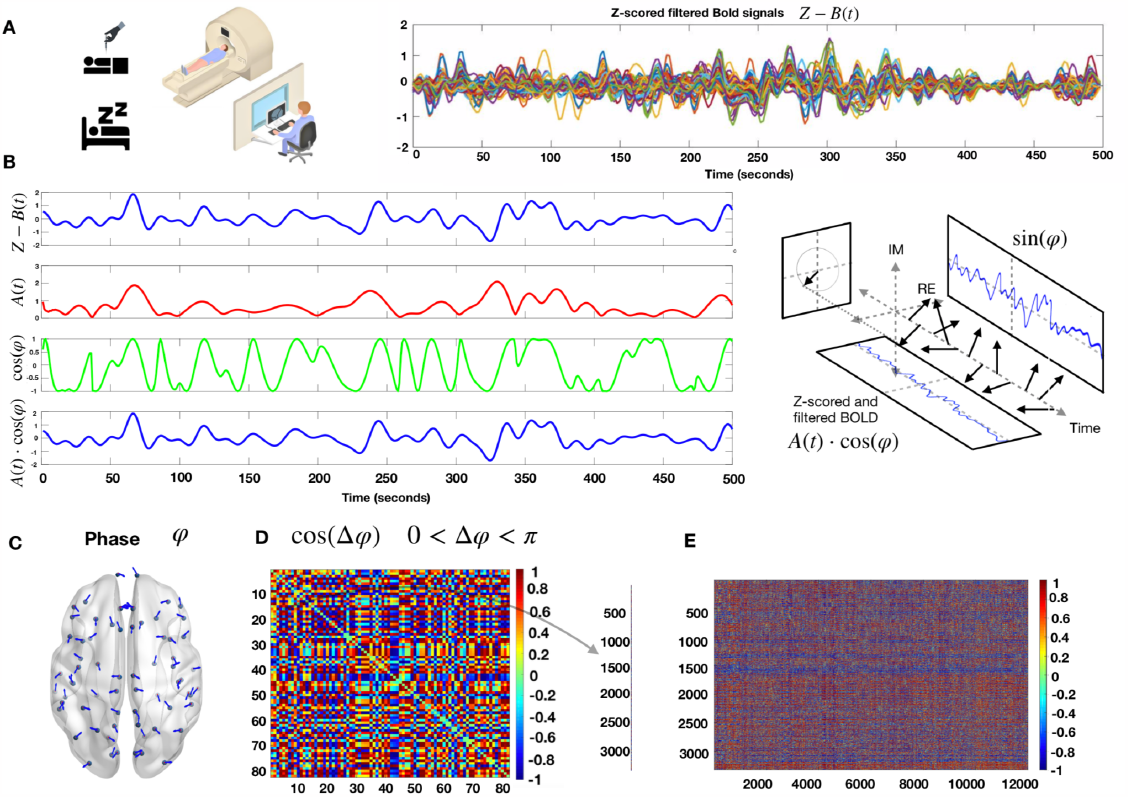
Phase-based dynamic functional patterns. **A)** Left- In the general anesthesia dataset, 16 healthy volunteers were scanned in three conditions (awake, G.A and recovery) and in the sleep database, 18 healthy volunteers were scanned in two conditions (awake and N3 sleep). Right) Example of Z-scored filtered BOLD time series for one participant of the anaesthesia database in the awake condition, 250 time points with a TR of 2s. **B)** Hilbert transform of the z-scored BOLD signal of one ROI into its time-varying amplitude A(t) (red) and the real part of the phase φ (green). In blue, we recover the original z-scored BOLD signal as A(t)cos(φ). **C)** Example of the phase of each brain region at one TR. **D)** Symmetric matrix of cosines of the phases differences between all pairs of brain regions. **E)** We concatenated the vectorized form of the triangular superior of the phase difference matrices for all TR’s for all participants, in all the conditions for both datasets separately.

### Instantaneous Phase

The analytic representation of a real-valued signal *x*(*t*) is a complex signal 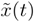 with the same Fourier transform as the real-valued signal, but defined only for positive frequencies. The analytic signal can be built from the real-valued signal using the Hilbert transform *H*:

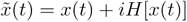

Where *i* is the imaginary unit. The main advantages of using the analytic signal are that, given some real-value data (for example BOLD signals), we can determine two functions of time to better access meaningful properties of the signal. We consider now a narrowband signal that can be written as an amplitude-modulated low-pass signal *A*(*t*) with carrier frequency expressed by *φ*(*t*) :

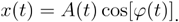

If the Fourier transforms of *A*(*t*) and cos[*φ*(*t*)] have separate supports, then the analytic signal of a narrowband signal can be rewritten as the product of two meaningful components

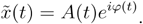

Where *A*(*t*) is the instantaneous amplitude and *φ*(*t*) is the instantaneous phase obtained from the Hilbert transform *H*[*x*(*t*)].

The narrower the bandwidth of the signal of interest, the better the Hilbert transform produces an analytic signal with a meaningful envelope and phase (Glerean et al. 2012). Adopting a band-pass filtered version of the BOLD time series improves the separation between the phase and envelope spectra.

The BOLD signals were first z-scored, and subsequently, we computed the Hilbert transform (Fig. 1B), to obtain the phase of each signal at each repetition time (TR), i.e. one time sample. Then, each instantaneous phase pattern was represented as a vector with N elements (here N = 68, since we used the Desikan Killiany atlas, comprising this number of ROIs), each element represents the projection of the phase (indicated by an arrow with an angle in Fig. 1C) of each brain area. Then, we built the matrix of phase coherence patterns (Fig. 1D) for each TR, as well as for each participant and condition.

### Phase-based dynamic functional coordination

To circumvent the issue of arbitrarily choosing the time-window and overlapping in capturing temporal oscillations employing a sliding-window methodology (Hindriks et al. 2016; Xie et al. 2019; Allen et al. 2014; Barttfeld, Uhrig, et al. 2015), the use of phase-based dynamic functional coordination was preferred (Alonso Martínez et al. 2020; Cabral et al. 2017; Vohryzek et al. 2020).

Analytic representations of signals were employed to derive a phase signal corresponding to the BOLD time series. We computed the instantaneous phase *φ*(*t*) of the BOLD signals across all ROI *n* ∈ {1, …, *N*} for each TR *t* ∈ {2, …, *T* − 1}, the first and last TR’s of each fMRI scan was excluded due to possible signal distortions induced by the Hilbert transform (Bracewell 2000).

The instantaneous phase was computed using Euler’s formula from the analytic signal, which was then “wrapped” within the range of −π to π, facilitating the calculation of inter-ROI phase differences. To obtain a whole-brain pattern of BOLD phase differences, the phase coherence between areas *k* and *j* at each time *t, PC*(*k,j,t*), was estimated using the pairwise phase coherence

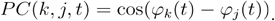

When areas *k* and *j* have synchronized BOLD signals at time *t*, phase coherence take value 1 and, when areas *k* and *j* in anti-phase at time *t* phase coherence is -1, all the rest of the cases lies in between the interval [−1, 1]. This computation was repeated for all subjects. For each subject, the resulting PC was a three-dimensional tensor with dimension *N* × *N* × *T*, where *N* is the number of regions in the parcellation considered (here 68 regions for the general anaesthesia and 90 regions for the deep sleep) and *T* the number of TR in each fMRI session.

### K-means clustering

To assess recurring coordination patterns among individuals in both datasets, a multistep methodology was employed. First, by taking advantage of the symmetry of the phase coherence matrices, we concatenated only the vectorized triangular superior parts of the phase coherence matrices for every time step together. Thus, the scanning sessions were transformed into matrices, wherein one dimension represented instantaneous phase differences (feature space) and the other dimension represented time (see Fig 1E) for all participants and all conditions. The resulting matrix was subjected to the k-means clustering algorithm utilising the L1 distance metric, as implemented in Python programming language (Scikit learn package). This resulted in a discrete set of *k* coordination patterns and their corresponding occurrence across time for each participant and each condition.

This process yielded *k* cluster centroids, which served as representatives of the recurring coordination patterns, accompanied by a label indicating the pattern to which each phase coherence matrix belongs according to the K-Means algorithm. The number of clusters was determined from a range from *k* = 3 to *k* = 10 (see Supplementary figures).

### Shannon Entropy

We make the histogram associated to the frequency of visits to each pattern for each participant in the different states. And for each of the normalized histogram of occurrence, for each participant, in each condition, and each value of *k*, we compute the Shannon entropy *S* as follows:

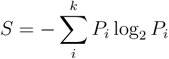

Where *P*_*i*_ is the probability (normalized histogram) of visit for each pattern for each participant in each brain state.

### Structure-function correlation

To investigate the dependence of brain dynamics on the state of consciousness, we defined a measure of similarity between functional and structural connectivity. We resorted to a widely used group estimate of human anatomical connectivity, measured with the noninvasive technique of Diffusion tensor imaging (DTI). We computed the linear correlation coefficient between the entries of both matrices (k-means centroids/anatomical connectome). This was performed for each phase-based coordination pattern obtained using the k-means algorithm, and was denoted SFC (see figures 2 and 4). Last, for each participant, we computed the linear slope coefficient of the relationship between the occurrence rate of each centroid and the corresponding centroid/anatomical correlation.

**Figure 2.**
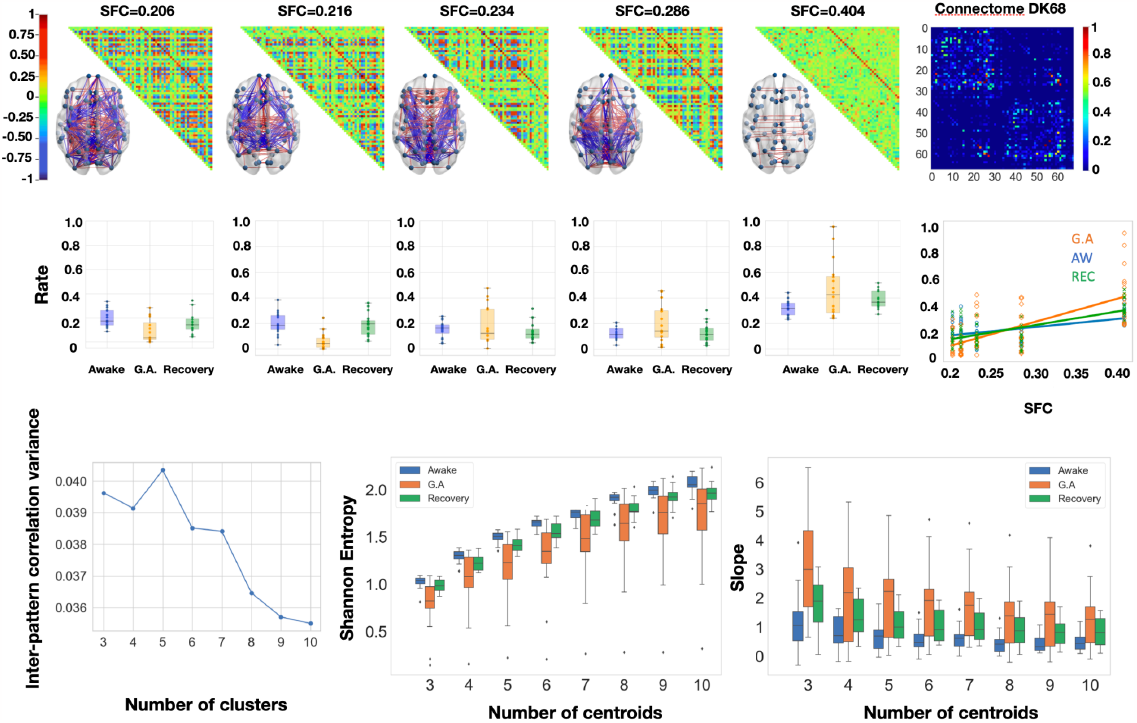
Anesthesia increases structure-function coupling and decreases Shannon entropy of distributions of occupancies of patterns. **A)** Matrix representation of the brain coordination patterns obtained for *k* = 5, which is the number of clusters that maximizes the inter-pattern correlation variance for 3 ≤ *k* ≤ 10, ordered by increasing SFC (from left to right). Only the strongest elements were depicted on the brain representation, in red the positive coherence > +0.4 and in blue the negative coherence < -0.4. Right: (normalized) connectome using the Desikan Killiany parcellation represented in its matrix form (68×68). **B)** Distribution of rates of occurrence for each of the above brain patterns across all 3 conditions. Right: Rate of the brain patterns as a function of their structure-function correlation (SFC). A linear regression is fitted with the mean rate of each brain pattern per condition. Statistical significance was evaluated with a paired t-test for the 3 pairs of conditions. **C)** Left: Inter-pattern correlation variance (metric used to determine the optimal number of brain patterns). Middle: Shannon Entropy of the distribution of occurrences of the *k* brain patterns 3 ≤ *k* ≤ 10 for the three conditions. Across all values of *k* analyzed, the same relative difference was found, under general anesthesia, the Shannon entropy is lower than in wakeful states. Right: Slopes of the linear regression for 3 ≤ *k* ≤ 10 computed for each participant, showing a tendency for anatomically dictated dynamics under general anesthesia. One-way repeated measure ANOVA was used to compute statistical differences between the conditions.

### Markov chains

For every time-point, the K-Means clustering assigns the index of the closest centroid to the empirical phase coherence matrix, resulting in a chain of indexes (integers, between 1 and *k*). We consider that sequence of centroids, or “patterns visited” for each participant at each time, as a sequence of random variables that jumps from one pattern to another pattern with a certain probability given by the empirical frequencies in which these events happen in the data. From this information we built a Markov Chain, as done previously (Demertzi et al. 2019).

More precisely, we counted the number of transitions between all pairs of patterns, excluding self-transitions which correspond to remaining in the same pattern, and normalized them to have a Markov transition matrix, where the sum of the rows is equal to one. We only include transitions of states of the same individual participant (see Figures 3 and 5), meaning the last state visited by a given patient and the first state visited by the next patient cannot be counted as a transition of state.

**Figure 3.**
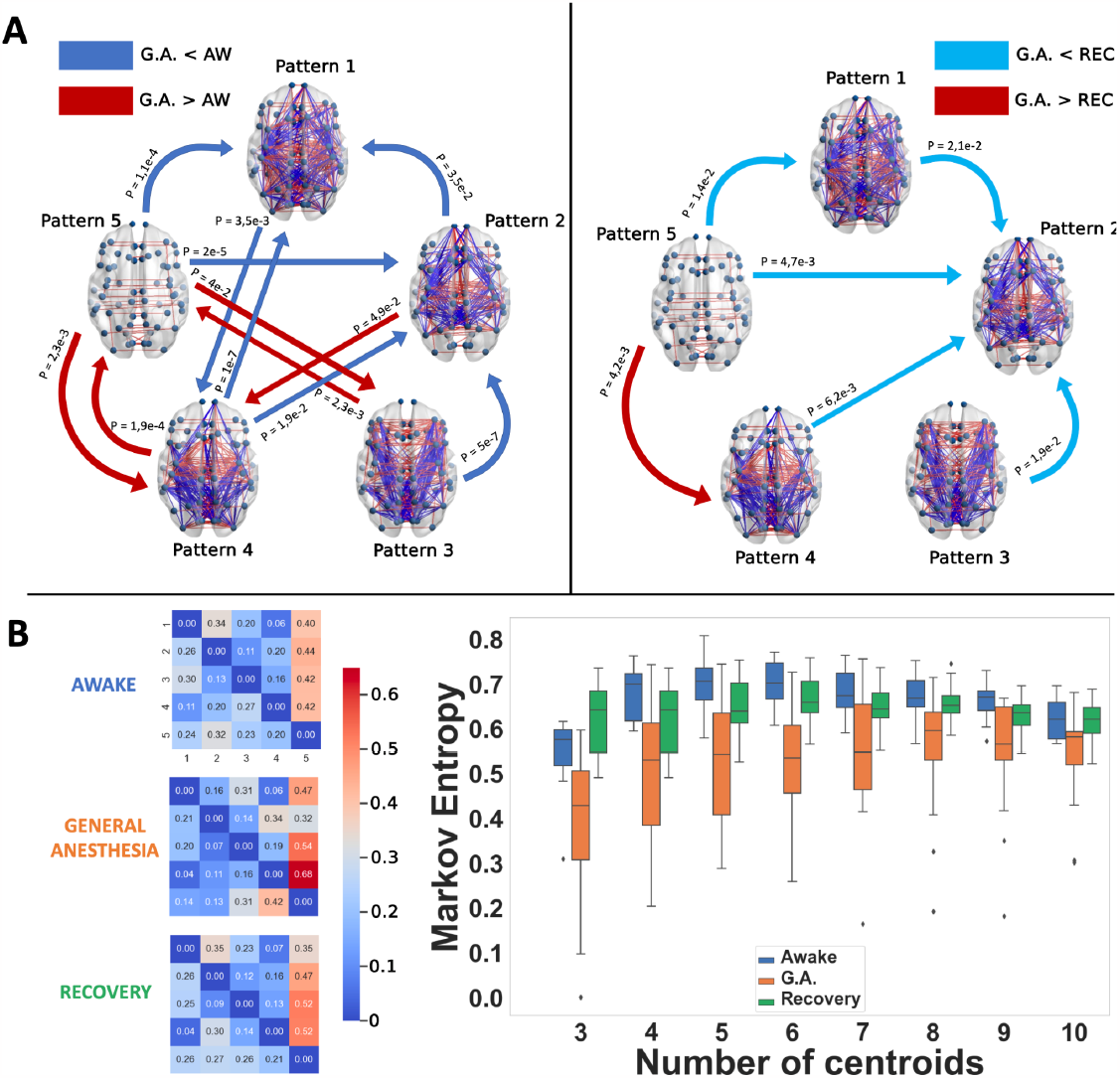
Propofol anesthesia decreases Markov entropy of transitions between patterns and produces robust transition differences with the conscious states, both awake and recovery. **A)** Brain network representations of the five different brain patterns and the statistical significance of the Markov transition probabilities using a two-sample paired t-test. Left: the awake condition is compared to general anesthesia (and vice-versa). Right: The general anesthesia condition is compared to the recovery condition. In blue are represented the transitions that were favoured in the wakeful conditions (awake and recovery), and in red the ones favoured in the unconscious condition. Only the transitions that passed the intermediate bootstrap check (CI=95%) were considered. Comparisons were made through paired t-tests. **B)** Left: Markov transition probabilities for the three conditions computed for *k* = 5. Right: Kolmgorov-Sinai entropy of the Markov chains obtained for 3 ≤ *k* ≤ 10. The Kolmgorov-Sinai entropy was consistently lower under general anesthesia. One-way repeated measure ANOVA was used to compute statistical differences between the conditions.

**Figure 4.**
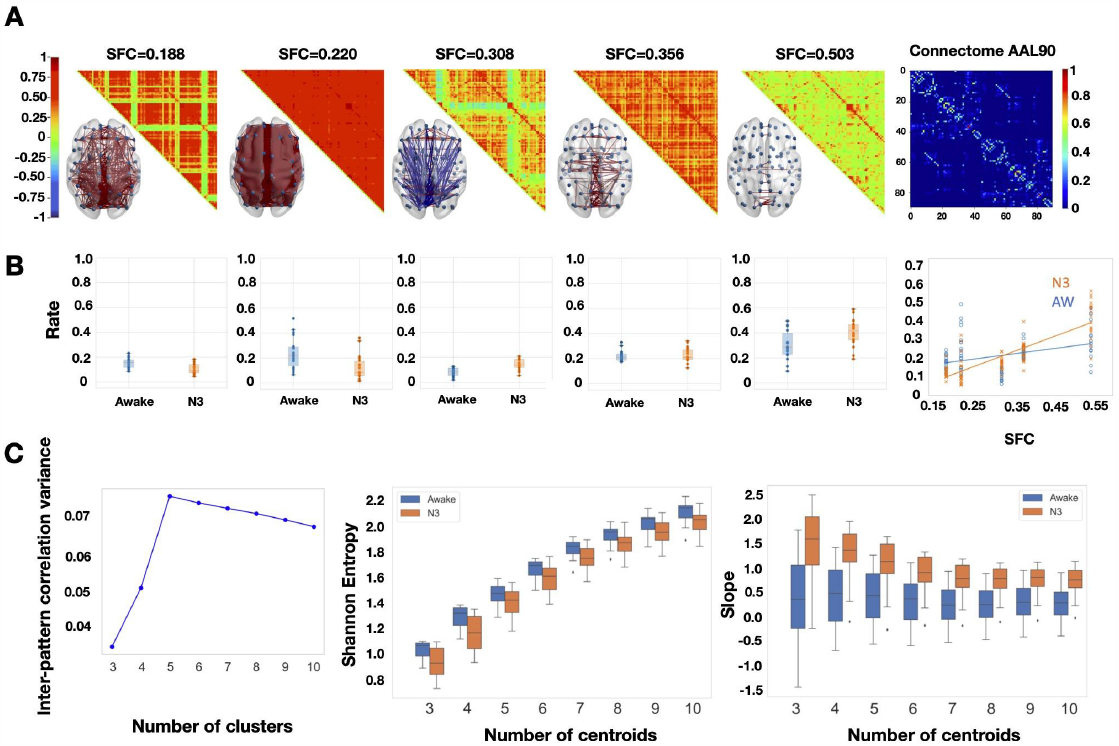
Deep sleep increases structure-function coupling and decreases Shannon entropy of distributions of occupancies of patterns. **A)** Matrix representation of the brain coordination patterns obtained for *k* = 5, which is the number of clusters that maximizes the inter-pattern correlation variance for 3 ≤ *k* ≤ 10, ordered by increasing SFC (from left to right). Only the strongest elements were depicted on the brain representation, in red the synchronizations > +0.4 and in blue anti-synchronizations < -0.4. Right: (normalized) connectome in the AAL90 parcellation (90×90). **B)** Distribution of rates of visit for each of the above brain patterns across the two conditions (Awake, N3 sleep). Paired t-test was used for comparison between conditions. Rate of the brain patterns as a function of their structure-function correlation (SFC). A linear regression is fitted with the mean rate of each brain pattern per condition, showing that, during deep sleep, the brain dynamics favor brain patterns with higher SFC unlike in the wakeful condition. **C)** Left: Inter-pattern correlation variance (metric used to determine the number of brain patterns). Middle: Shannon Entropy of the distribution of rates *k* of the brain patterns 3 ≤ *k* ≤ 10 for two conditions. Across all values of *k*, the same relative difference is found, during profound sleep, the Shannon entropy is lower than in wakeful states. Right: Slopes of the linear regressions for 3 ≤ *k* ≤ 10, computed for each participant, showing a tendency for anatomically dictated dynamics in deep sleep. Paired t-test was used for comparaison between conditions.

**Figure 5.**
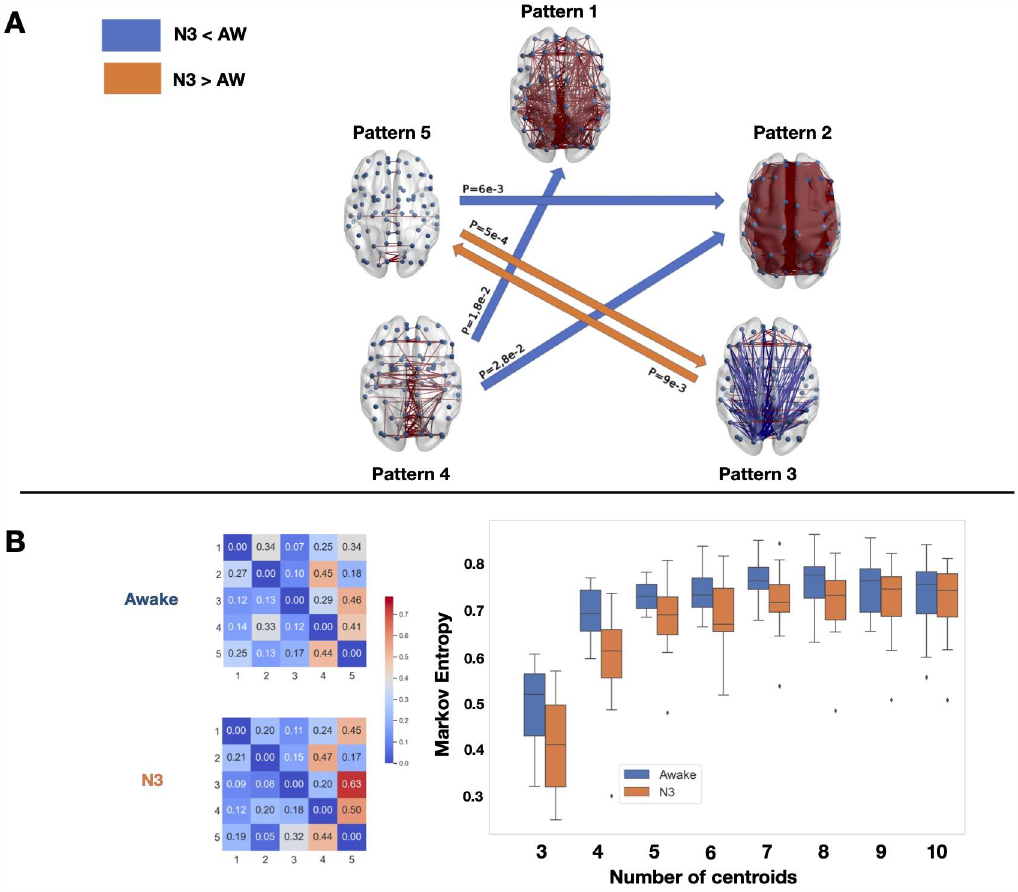
Deep Sleep decrease Markov entropy of transitions between patterns and produce robust transition differences with the conscious states awake and recovery. **A)** Brain network representations of the four different brain patterns and the statistical significance of the Markov transition probabilities using a t-test. The awake condition is compared to sleep (and vice-versa). In blue are represented the transitions that were favored in the awake condition, and in orange the ones favored in the deep sleep condition (N3). Again, only the transitions that passed the intermediate bootstrap check (CI=95%) are showed. **B)** On the left are the Markov transition probabilities for the two conditions computed for *k* = 5. On the right is the Kolmgorov-Sinai entropy of the Markov chains obtained for 3 ≤ *k* ≤ 10. The Kolmgorov-Sinai entropy was consistently lower under general anesthesia.

In order to generalize the notion of uncertainty of a probability distribution given by its Shannon entropy, here we quantify the degree of randomness or uncertainty there is in a sequence of random variables characterised by a discrete Markov Chain. For each participant from their Markov transition matrix, we obtained the stationary distribution *μ* represented as a row vector whose entries are probabilities, where *μ*_*i*_ corresponds to the stationary probability of observing pattern *i* (i.e, over the long run, independent of the starting pattern, the proportion of time the chain spends in state *i* is approximately *μ*_*i*_ for all *i*). The stationary probability is computed as follows: *μ***P** = *μ*, where **P** is the Markov transition matrix.

Then, we calculate the entropy rate, or Kolmogorov-Sinai entropy, of the Markov chains as follows:

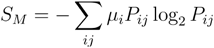

where *P*_*ij*_ corresponds to the transition probability from pattern *i* to pattern *j*.

### Statistical methods

To assess the statistical significance of the differences in, transition probabilities, slope’s coefficients, and Shannon entropy, between conditions, we performed two paired t-test with Bonferroni correction between the awake and general anesthesia conditions, and between the recovery and general anesthesia ones, and a repeated measure ANOVA. Similarly, we performed one paired t-test to compare the N3-associated measurements and the awake ones. Then, in order to ensure that the transition probabilities differences between conditions were indeed due to the specificity of the brain conditions, we resorted to a bootstrap method for each subject. For every individual, we randomly shuffled the indexes from the chain of states 10.000 times and computed as many transition matrices, thus breaking any temporal dependencies of the transitions between specific states. We then verified if the actual transitions did not belong to a confidence interval, successively designed by a confidence level of 90%, 95%, and 99%, defined by the shuffled transitions matrices.

## Results

We investigated a dataset of anaesthesia with the intravenous agent propofol, where the BOLD signals of 16 participants during three conditions (awake, general anesthesia and recovery) and a dataset of deep sleep consisting of 18 participants in two conditions (awake and N3 sleep). Both datasets were obtained following the procedures indicated in detail in the section “fMRI acquisition, experimental design and processing”.

Our phase-based analysis consistently unveiled discernible cerebral patterns of brain dynamics, demonstrating robustness against the choice of the number of clusters in the k-means algorithm (see Supplementary Figure S1 and S2). Notably, the brain pattern possessing the lowest correlation with the connectome exhibited higher occurrence during the awake state in comparison to participants under general anesthesia (Fig. 2) and N3 sleep (Fig. 4). In stark contrast, a pattern indicative of heightened structure-function correlation was more frequently observed in participants under general anesthesia and deep sleep, as opposed to those in the awake state (see Figure 2B and Figure 4B).

To illustrate our results we choose the number of patterns that maximized the inter-pattern correlation variance as in previous studies (Demertzi et al. 2019), that turns out to be five for the general anesthesia dataset and four for deep sleep dataset. However, as shown throughout this article our results are robust independently of the choice of the number of clusters from three to ten. We show that the slope of structure-function correlation with respect to the rate of occurrence of the patterns is higher when consciousness is suppressed either by general anesthesia or deep sleep (Fig 2B right and 4B right) independently of the choice of the number of centroids (patterns). We also show that the Shannon entropy computed from the normalized histograms (probability distributions) of occurrences of each brain pattern for each condition is higher in the awake and recovery conditions with respect to general anesthesia (Fig 2C middle and 4C middle). A similar behavior is observed in the deep sleep dataset, where the Shannon entropy is higher in the awake state with respect to the N3 stage. In both datasets the result is robust to the choice of the number of centroids (see Figures 2,3,4 and 5).

Finally, we computed in both datasets the frequency of transitions between patterns. We selected the statistically significant pattern transitions obtained by comparing pairs of conditions for general anesthesia and deep sleep (Fig 3 and 5). From the frequency of transitions between patterns we estimated Markov chains for each condition, from which we computed the Markov Entropy. We observed higher Markov entropy for the conscious states (awake and recovery) with respect to general anesthesia, and a similar result contrasting the awake state versus N3. Both results generalize well with different choices of the number of centroids.

## Discussion

The present study delves into the generalization of results obtained previously (Demertzi et al. 2019) about the dynamics of the human brain and how they are altered in states of loss of consciousness, by employing two complementary datasets of humans under general anesthesia and N3 sleep. The ability to employ the same method effectively across independent datasets in different scenarios of low consciousness, emphasizes its reliability and underscores its potential to yield consistent outcomes in other datasets. Such cross-applicability is indicative of the robustness of the approach and the generalizability of the results. Specifically, loss of consciousness - whether due to spontaneous sleep, or induced by general anaesthesia - appears to reduce the repertoire of dynamical states that the brain visits, with increased prevalence of the most structurally-coupled pattern.

Our results demonstrate that sustaining rich brain dynamics is essential for consciousness and can serve as a biomarker for consciousness. While the awake state exhibits a richer dynamic exploration of the functional repertoire as exhibited by the Shannon entropy analyzing the histograms (Fig 2C and 4C) and the Markov entropy for the Markov chains (fig 3B and 5B), the dynamic exploration is consistently reduced under anesthesia and N3 sleep for all the different choices for the number of clusters. This result is well aligned with the entropic brain hypothesis (Carhart-Harris et al. 2014; Carhart-Harris 2018; Herzog et al. 2023). This diversity in state exploration is also related to recent results in the mouse using calcium imaging under different drugs of general anaesthesia, which show that under anaesthesia, the brain explore less states than the awake brain (Filipchuk et al. 2022), and recent results of the structure-function interdependence of the macaque brain under loss of consciousness induced by three different anaesthetics (sevoflurane, propofol, ketamine) and restoration of consciousness by deep brain stimulation (DBS) (Luppi et al. 2023). Furthermore, the more prevalent functional connectivity patterns during anesthesia correlate with the anatomical connectivity, consistent with previous findings. As in previous studies (Demertzi et al. 2019), here we find that the slope of occurrence probability versus structure-function correlation increases in the states of low consciousness. Additionally, here we show that those earlier results generalize to general anesthesia and N3 sleep and different values of brain patterns.

A notable feature of the phase based methodology is that it does not need meticulous adjustment of the width of sliding windows. This autonomy from fine-tuning aspects diminishes the potential for bias introduced by window length selection, highlighting again the robustness of the methodology.

The sporadic emergence of patterns more present in states of low consciousness during periods of conscious wakefulness, and vice versa, raises pertinent questions about the boundaries between these brain states. The co-occurrence of seemingly contrasting states within the same individual challenges conventional notions of discrete consciousness states. This observation requires further investigation into the underlying mechanisms that give rise to these sporadic occurrences and offers a unique perspective on the dynamic nature of consciousness (Andrillon et al. 2021, 2019; Mortaheb et al. 2022; Kawagoe, Onoda, and Yamaguchi 2019). The alterations induced by propofol in the body and the brain is not only reduced to consciousness, as elucidated by (Barttfeld, Bekinschtein, et al. 2015). Discrimination between changes stemming directly from the loss of consciousness and those arising from ancillary effects of propofol on cerebral processes remains challenging. A comparable challenge is encountered in the realm of sleep research, as underscored by (Siclari et al. 2017), wherein deep sleep is acknowledged as more than mere unconsciousness. Additionally, diminished vigilance is observed independently of attenuated awareness (Oken, Salinsky, and Elsas 2006).

The question arises concerning the generalizability of the findings through alternative brain recording techniques, bypassing the reliance on functional Magnetic Resonance Imaging (fMRI). The prospect of employing different data sources, such as electroencephalography (EEG) or magnetoencephalography (MEG), to substantiate the identified consciousness states introduces an avenue for expanding the horizon of the current methodology, and enhances the robustness of the method and the findings (Vidaurre et al. 2018, 2016; Baker et al. 2014; Sitt et al. 2014). Generalisations of the present method to EEG are under development (Della Bella et al. 2022), and may be of particular relevance, for example, for the study of unconsciousness induced by epileptic seizures. Alternatively, other species under general anaesthesia can be studied as has been done in the past with the sliding window technique (Barttfeld, Uhrig, et al. 2015; Uhrig et al. 2018) Envisioning a broader context, the study prompts consideration of consciousness-altering scenarios beyond those encountered within wakefulness or unconscious states.

However, it is important to acknowledge the limitations inherent in the study. Important gaps remain in understanding how these dynamics are influenced by brain structure and anesthetic agents. For example another widely used anaesthetic, sevoflurane, has been shown to resemble propofol in terms of its effects on brain dynamics, both in humans (Golkowski et al. 2019; Luppi et al. 2019; Luppi, Golkowski, et al. 2021) and macaques (Uhrig et al. 2018; Signorelli et al. 2021). Another important limitation is that we do not know which changes are a consequence of propofol reducing consciousness, and which appear because propofol does other things in the brain (Barttfeld, Bekinschtein, et al. 2015). The same problem is present for sleep, since deep sleep is not only loss of consciousness (consciousness may be present as shown by Siclari, Tononi, et al. (Siclari et al. 2017)), and reduced vigilance occurs independently of reduced awareness (Oken, Salinsky, and Elsas 2006). The generality of the method is also partly a weakness, because as can be seen in figures 2 and 4, the patterns of general anesthesia and deep sleep are quite different. This may be due to the fact that they are obtained from data obtained by different research groups using different data acquisition parameters, with different pre-processing protocols and different participants. Despite these limitations, the method appears to be robust and generalisable; the question then remains: how broad is the generalisability of the present findings?

In conclusion, the present study highlights the generalization and robustness of previous results employing a uniform methodology across varying states exhibited by different cohorts, in different consciousness states. By leveraging the complementary nature of our results, we provide a comprehensive characterization of the dynamic features of brain networks and how consciousness reshapes the dynamics of the human brain and its relation with the connectome. This study contributes to the broader discourse on consciousness and methodology, paving the way for future investigations in the relationship of consciousness and dynamical brain patterns and to identify potential markers that may be utilized in the future to distinguish between conscious and unconscious states.

### fMRI acquisition, experimental design and processing

#### Anaesthesia data: Recruitment

The propofol data employed in this study have been published before (Luppi et al. 2019; Varley et al. 2020; Naci et al. 2018; Kandeepan et al. 2020). For clarity and consistency of reporting, where applicable we use the same wording as our previous studies. The propofol data were collected between May and November 2014 at the Robarts Research Institute in London, Ontario (Canada) (Luppi et al. 2019). The study received ethical approval from the Health Sciences Research Ethics Board and Psychology Research Ethics Board of Western University (Ontario, Canada). Healthy volunteers (n = 19) were recruited (18–40 years; 13 males). Volunteers were right-handed, native English speakers, and had no history of neurological disorders. In accordance with relevant ethical guidelines, each volunteer provided written informed consent, and received monetary compensation for their time. Due to equipment malfunction or physiological impediments to anaesthesia in the scanner, data from n = 3 participants (1 male) were excluded from analyses, leaving a total n = 16 for analysis.

#### Anaesthesia data: Procedure

Resting-state fMRI data were acquired at different propofol levels: no sedation (Awake), and Deep anaesthesia (corresponding to Ramsay score of 5). As previously reported (Luppi et al. 2019), for each condition fMRI acquisition began after two anaesthesiologists and one anaesthesia nurse independently assessed Ramsay level in the scanning room. The anaesthesiologists and the anaesthesia nurse could not be blinded to experimental condition, since part of their role involved determining the participants’ level of anaesthesia. Note that the Ramsay score is designed for critical care patients, and therefore participants did not receive a score during the Awake condition before propofol administration: rather, they were required to be fully awake, alert and communicating appropriately. To provide a further, independent evaluation of participants’ level of responsiveness, they were asked to perform two tasks: a test of verbal memory recall, and a computer-based auditory target-detection task. Wakefulness was also monitored using an infrared camera placed inside the scanner.

Propofol (a potent agonist of inhibitory GABA-A receptors (Yip et al. 2013; Jurd et al. 2003) was administered intravenously using an AS50 auto syringe infusion pump (Baxter Healthcare, Singapore); an effect-site/plasma steering algorithm combined with the computer-controlled infusion pump was used to achieve step-wise sedation increments, followed by manual adjustments as required to reach the desired target concentrations of propofol according to the TIVA Trainer (European Society for Intravenous Aneaesthesia, eurosiva.eu) pharmacokinetic simulation program. This software also specified the blood concentrations of propofol, following the Marsh 3-compartment model, which were used as targets for the pharmacokinetic model providing target-controlled infusion. After an initial propofol target effect-site concentration of 0.6 μg mL−1, concentration was gradually increased by increments of 0.3 μg mL1, and Ramsay score was assessed after each increment: a further increment occurred if the Ramsay score was lower than 5. The mean estimated effect-site and plasma propofol concentrations were kept stable by the pharmacokinetic model delivered via the TIVA Trainer infusion pump. Ramsay level 5 was achieved when participants stopped responding to verbal commands, were unable to engage in conversation, and were rousable only to physical stimulation. Once both anaesthesiologists and the anaesthesia nurse all agreed that Ramsay sedation level 5 had been reached, and participants stopped responding to both tasks, data acquisition was initiated. The mean estimated effect-site propofol concentration was 2.48 (1.82–3.14) μg mL−1, and the mean estimated plasma propofol concentration was 2.68 (1.92–3.44) μg mL−1. Mean total mass of propofol administered was 486.58 (373.30–599.86) mg. These values of variability are typical for the pharmacokinetics and pharmacodynamics of propofol. Oxygen was titrated to maintain SpO2 above 96%. At Ramsay 5 level, participants remained capable of spontaneous cardiovascular function and ventilation. However, the sedation procedure did not take place in a hospital setting; therefore, intubation during scanning could not be used to ensure airway security during scanning. Consequently, although two anaesthesiologists closely monitored each participant, scanner time was minimised to ensure return to normal breathing following deep sedation. No state changes or movement were noted during the deep sedation scanning for any of the participants included in the study.

#### Anaesthesia data: Design

As previously reported (Luppi et al. 2019), once in the scanner participants were instructed to relax with closed eyes, without falling asleep. Resting-state functional MRI in the absence of any tasks was acquired for 8 min for each participant. A further scan was also acquired during auditory presentation of a plot-driven story through headphones (5 min long). Participants were instructed to listen while keeping their eyes closed. The present analysis focuses on the resting-state data only; the story scan data have been published separately (Kandeepan et al. 2020).

#### Anaesthesia data: fMRI data acquisition

As previously reported (Luppi et al. 2019), MRI scanning was performed using a 3-Tesla Siemens Tim Trio scanner (32-channel coil), and 256 functional volumes (echo-planar images, EPI) were collected from each participant, with the following parameters: slices = 33, with 25% inter-slice gap; resolution = 3 mm isotropic; TR = 2000 ms; TE = 30 ms; flip angle = 75 degrees; matrix size = 64 × 64. The order of acquisition was interleaved, bottom-up. Anatomical scanning was also performed, acquiring a high-resolution T1-weighted volume (32-channel coil, 1 mm isotropic voxel size) with a 3D MPRAGE sequence, using the following parameters: TA = 5 min, TE = 4.25 ms, 240 × 256 matrix size, 9 degrees flip angle (Luppi et al. 2019).

#### Sleep data: Recruitment

A total of 63 healthy subjects (36 females, mean SD, 23.4 3.3 years) were selected from a data set previously described in a sleep-related study by Tagliazucchi and Laufs (Tagliazucchi and Laufs 2014). Participants entered the scanner at 7 PM and were asked to relax, close their eyes, and not fight the sleep onset. A total of 52 minutes of resting state activity were measured with a simultaneous combination of EEG and fMRI. According to the rules of the American Academy of Sleep Medicine, the polysomnography signals (including the scalp potentials measured with EEG) determine the classification of data into four stages (wakefulness, N1, N2, and N3 sleep).

We selected 18 subjects with contiguous resting-state time series of at least 200 volumes to perform our analysis. The local ethics committee approves the experimental protocol (Goethe-Universität Frankfurt, Germany, protocol number: 305/07), and written informed consent was asked to all participants before the experiment. The study was conducted according to the Helsinki Declaration on ethical research.

#### Sleep data: MRI data acquisition

MRI images were acquired on a 3-Tesla Siemens Trio scanner (Erlangen, Germany) and fMRI acquisition parameters were 1505 volumes of T2-weighted echo planar images, TR/TE = 2080ms/30ms, matrix 64×64, voxel size 3×3×3 *mm*^3^, distance factor 50%; FOV 192 *mm*^2^. An optimized polysomnographic setting was employed (chin and tibial EMG, ECG, EOG recorded bipolarly [sampling rate 5 kHz, low pass filter 1 kHz] with 30 EEG channels recorded with FCz as the reference [sampling rate 5 kHz, low pass filter 250 Hz]. Pulse oximetry and respiration were recorded via sensors from the Trio [sampling rate 50 Hz]) and MR scanner-compatible devices (BrainAmp MR+, BrainAmpExG; Brain Products, Gilching, Germany), facilitating sleep scoring during fMRI acquisition.

#### Sleep data: Brain parcellation AAL 90 to extract BOLD time series and filtering

To extract the time series of BOLD signals from each participant in a coarse parcellation, we used the AAL90 parcellation with 90 brain areas anatomically defined in [25]. BOLD signals (empirical or simulated) were filtered with a Butterworth (order 2) band-pass filter in the 0.01-0.1 Hz frequency range.

## Data availability statement

The dataset of general anesthesia is openly available and can be found in the following link https://openneuro.org/datasets/ds003171/versions/1.0.0

The dataset of sleep is available on request from

## Code availability

All analyses were carried out using custom scripts written in Python and openly available in the Github repository https://github.com/Krigsa/phase_coherence_kmeans.

## Supplementary Information

### Supplementary figures

**Figure S1.**
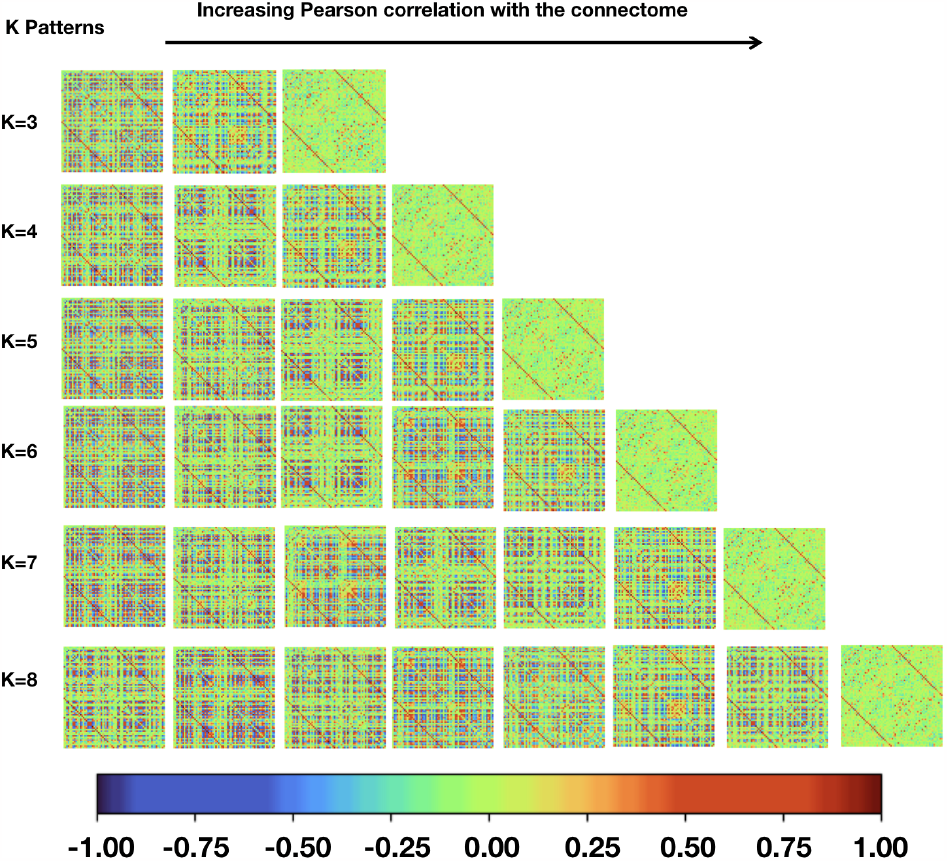
Dynamic functional coordination patterns. From propofol general anesthesia dataset using K = 3.. 8 number of clusters in the k-means algorithm. For each value of K patterns are ordered based on their similarity to underlying anatomical connectivity, from the least (left) to the most similar (right).

**Figure S2.**
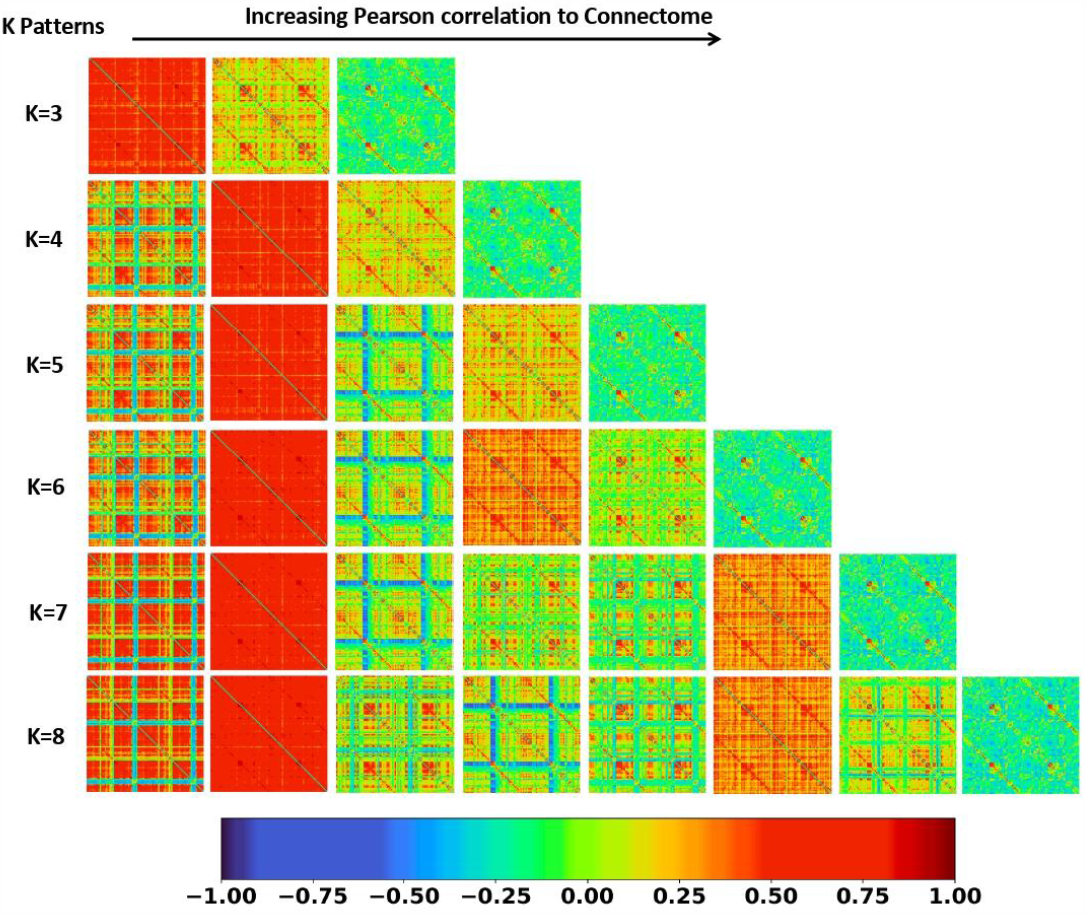
Dynamic functional coordination patterns. From the dataset of sleep using K = 3.. 8 number of clusters in the k-means algorithm. For each value of K patterns are ordered based on their similarity to underlying anatomical connectivity, from the least (left) to the most similar (right).

